# Fine-scale adaptive divergence of *Aedes aegypti* in heterogeneous landscapes and among climatic conditions in Metropolitan Manila, Philippines

**DOI:** 10.1101/2023.04.12.536525

**Authors:** Atikah Fitria Muharromah, Thaddeus M. Carvajal, Maria Angenica F. Regilme, Kozo Watanabe

**Affiliations:** Center for Marine Environmental Studies (CMES), Ehime University, Bunkyo-cho 3, Matsuyama, Ehime 7908577, Japan; Graduate School of Science and Engineering, Ehime University, Bunkyo-cho 3, Matsuyama, Ehime 7908577, Japan; Department of Tropical Biology, Faculty of Biology, Universitas Gadjah Mada, Yogyakarta 55281, Indonesia; Biological Control Research Unit, Center for Natural Sciences and Environmental Research, De La Salle University, 2401 Taft Avenue, Manila 1004, Philippines

**Author notes:** Corresponding author (KW).

## Abstract

The adaptive divergence of *Aedes aegypti* populations to heterogeneous environments may be a driving force behind the recent expansion of their habitat distribution and outbreaks of dengue disease in urbanized areas. In this study, we investigated the population genomics of *Ae. aegypti* at a regional scale in Metropolitan Manila, Philippines using double digestion restriction-site association DNA sequencing (ddRAD-Seq). Specifically, we used a Pool-Seq approach to generate a high number of single nucleotide polymorphisms (SNPs), which were used to determine local adaptation and population structure. We detected 65,473 SNPs in 217 *Ae. aegypti* individuals from 14 populations with 76 non-neutral SNP loci. Additionally, 57 of these non-neutral SNP loci were associated with 8 landscape variables (e.g., open space, forest, etc) and 4 climate variables (e.g., air temperature, humidity, etc). Furthermore, the percentage of the area of landscape variables, such as forest, parks and recreation, air temperature, man-made building, and open space per local population was frequently associated with non-neutral SNP loci. Most non-neutral SNP loci formed four clusters that were in linkage disequilibrium with each other in physical proximity on the chromosome and were associated with a common environmental variable. Male and female populations exhibited contrasting spatial divergence, i.e., males exhibited greater divergence, likely reflecting their different dispersal abilities. In comparative analysis of the same *Ae. aegypti* individuals, the pairwise *F*_ST_ values of 11 microsatellite markers were lower than those of neutral SNP loci, indicating that the neutral SNP loci generated via ddRAD-Seq were more sensitive in terms of detecting genetic differences between populations at fine-spatial scales. Overall, this study demonstrates the utility of ddRAD-Seq for examining genetic differences in *Ae. aegypti* populations, and our data on mosquito dispersal at a regional spatial scale could inform vector control programs.

**Author Summary:** The population expansion of dengue vector, *Aedes aegypti* mosquitoes is one of the factors that may promote the outbreak of the diseases. Understanding the population genomics of *Ae.aegypti* may contribute to better knowledge about mosquito expansion and how they can adapt to the change in environment. In this study, we used pool-based ddRAD-Seq (Double Digest Restriction site Association DNA Sequencing) to generate SNPs that occur between the *Ae.aegypti* populations in Metropolitan Manila, Philippines. We found that non-neutral SNP loci are frequently associated with landscape variables compared to climatic variables. Landscape variables such as forest, park and recreation, air temperature, man-made building and open space are more frequently associated with non-neutral SNPs loci. Those landscape variables may relate to the mosquito’s fitness, therefore, induce the adaptive divergence within *Ae.aegypti* population. We also found male and female populations are exhibiting a contrast spatial divergence by using neutral SNP loci. In addition, neutral SNPs loci showed higher resolution in population structuring than microsatellite markers using the same individuals.

## Introduction

*Aedes aegypti* is an important vector of mosquito-borne diseases, including dengue disease [1,2]. In recent decades, dengue disease cases have increased in urbanized areas [3], possibly owing to the recent habitat expansion of *Ae. aegypti* [4,5], which suggests that *Ae. aegypti* possess adaptive genes that function in emerging urban environments (e.g., human settlements). In addition to high adaptability to urban environments, the high dispersal ability of *Ae. aegypti* may contribute to their habitat expansion [1,6]. Understanding the ecology of vectors with respect to their environmental adaptation and dispersal ability may allow us to predict the expansion of their habitat distribution under changing environmental conditions, such as changing landscape and weather conditions [6,7]. Moreover, we can obtain this understanding through population genomics and landscape genomics approaches [8,9].

Double digestion restriction-site association DNA sequencing (ddRAD-Seq) facilitates genomic analysis by generating a high number of single nucleotide polymorphisms (SNPs) [10,11]. Among the many SNP loci, the few loci affected by directional selection should exhibit greater genetic differentiation than the neutral loci comprising the majority of the genome, whereas the few loci subject to balancing selection should exhibit lower genetic differentiation. These “outlier” loci can be identified as non-neutral loci through statistical methods. The environmental factors that cause natural selection can be estimated based on the correlation between non-neutral loci and environmental variables. Additionally, non-neutral loci in physical proximity on a chromosome often share similar effects of natural selection owing to the reduced recombination occurring at that part of the genome, leading to linkage disequilibrium. Therefore, if many non-neutral loci associated with environmental variables are identified, it becomes possible to analyze their linkage relationships.

Few studies have focused on the adaptive divergence of *Ae. aegypti* along an environmental gradient [12,13]. Sherpa et al. [12] identified potential adaptive loci associated with human density and/or insecticide resistance at a continental scale, i.e., Africa and the Caribbean. A national-scale study in Panama revealed that *Ae. aegypti* populations were undergoing adaptive divergence along environmental gradients of temperature and vegetation [13]. However, adaptive divergence has not previously been examined at a regional scale, e.g., within a city, in which spatial genetic variance and environmental heterogeneity are usually low.

Neutral loci have been studied extensively to understand neutral evolutionary processes, including migration and genetic drift. Microsatellite markers, which are neutral markers [14], are widely used in population genetics analysis owing to their high levels of mutagenicity and polymorphism. However, a limited number of microsatellite markers might limit the resolution power in respect of resolving the genetic structure among evolutionarily closely related populations co-occurring at a fine-spatial scale [15]. SNP loci generated in abundance through next-generation sequencing (NGS) via ddRAD-Seq are preferably used in population genetics because they allow the clear detection of population genetic structure, even at a fine-spatial scale [16]. Rašić et al. [19] compared the ability of microsatellite markers and several SNP loci found through ddRAD-Seq to detect genetic differentiation among populations at continental [19] and city spatial scales [23,24], finding that the SNP loci could detect more distinct genetic differentiation among populations than the microsatellite markers. An additional advantage of ddRAD-Seq over microsatellite markers is obtaining neutral and non-neutral loci that can be used for studying population structure and adaptive divergence simultaneously. However, one limitation of ddRAD-Seq is its high cost, which may preclude the analysis of a large number of individuals. Nevertheless, larger sample sizes in a population are better for accurately estimating allele frequencies in the population [17]. Accordingly, Pool-Seq, a sequencing strategy that greatly reduces the cost and time of ddRAD-Seq by pooling multiple individual samples, has been developed [18]. Pool-Seq can estimate the gene frequencies of many populations relatively inexpensively because many individuals are available per population.

Population genetics studies on *Ae. aegypti* have mainly focused on female mosquitoes, which transmit diseases. However, the population structure of male *Ae. aegypti* populations must also be explored in respect of its function in mosquito control strategies, e.g., the *Wolbachia*–*Aedes* suppression strategy, and the release of sterile male mosquitoes into populations [28]. In Metropolitan Manila, Philippines, Carvajal et al. [29] separated female and male populations using microsatellite markers, thereby revealing their different dispersal patterns. At the fine-spatial scale, females and males in the same population tend to be highly genetically similar and difficult to separate. Therefore, determining the population structures of female and male *Ae. aegypti* at a fine-spatial scale with confidence requires many neutral markers, e.g., SNP markers. However, the population genomics of female and male *Ae. aegypti* populations, including their adaptive divergence and population structure on a fine-spatial scale, have not been studied.

The present study aims to determine the population genomic structure of *Ae. aegypti* mosquitoes at a regional spatial scale, i.e., in Metropolitan Manila. The specific objectives of the study are as follows: (1) to identify the adaptive divergence of *Ae. aegypti* along environmental gradients of climatic and/or landscape factors across the regional scale; (2) to examine linkage among non-neutral loci potentially undergoing natural selection in the *Ae. aegypti* genome; (3) to compare the population divergence levels of female and male *Ae. aegypti*; and (4) to determine whether a number of SNP loci detected via ddRAD-Seq or microsatellite markers are more capable of detecting genetic differentiation among populations on a regional scale. Regarding these aims, we successfully used pooled ddRAD-Seq to detect adaptive divergence among *Ae. aegypti* populations along environmental gradients at a relatively fine-spatial scale in Metropolitan Manila, and we determined dispersal patterns among local populations as well as their sex differences.

## Methods

### Study area

We performed ddRAD-Seq analysis using DNA sampled from 217 *Ae. aegypti* individuals collected from Metropolitan Manila, which had previously been used in two population genetic studies using 11 microsatellite markers [26,27]. Of the 217 individuals, 165 were collected from 82 households distributed throughout Metropolitan Manila [26], whereas 52 were collected intensively from 39 households distributed in a small area (0.048 km^2^) in Manila, Metropolitan Manila [27]. The samples were collected using a UV light trap (MosquitoTrap, Jocanima Corporation, Las Pinas City, Philippines) from May 2014 to January 2015 [26] and from September to October 2017 [27]. The individual insects were identified at species level using the pictorial keys of Rueda et al. [30]. The same DNA samples previously collected from Metropolitan Manila [26, 27] were used in the present study to allow a comparison between the previously generated microsatellite marker data and the SNPs marker data generated in the current study.

The present study considered 14 *Ae. aegypti* populations, including 7 male and 7 female populations, from 5 locations in Metropolitan Manila: North, West, East, Central, South regions studied by Carvajal et al. [26] and two additional locations in a small spatial area of Manila, namely Manila North and Manila South, studied by Regilme et al. [27] (Fig. 1). In total, 153 households were considered in the present study. The number of households per male and female population was 7–15 and 8–16 (mean: 10.1 and 11.7 households), respectively (S1 Table). Because the households per population were scattered, we calculated the geographical midpoint of nearby households using ArcGIS 10.2 (ESRI, Redlands, California) and determined this as a population. The total number of male and female *Ae. aegypti* samples was 93 and 124 individuals, with 7–20 and 12–28 individuals per population, respectively (S1 Table).

**Fig 1.**
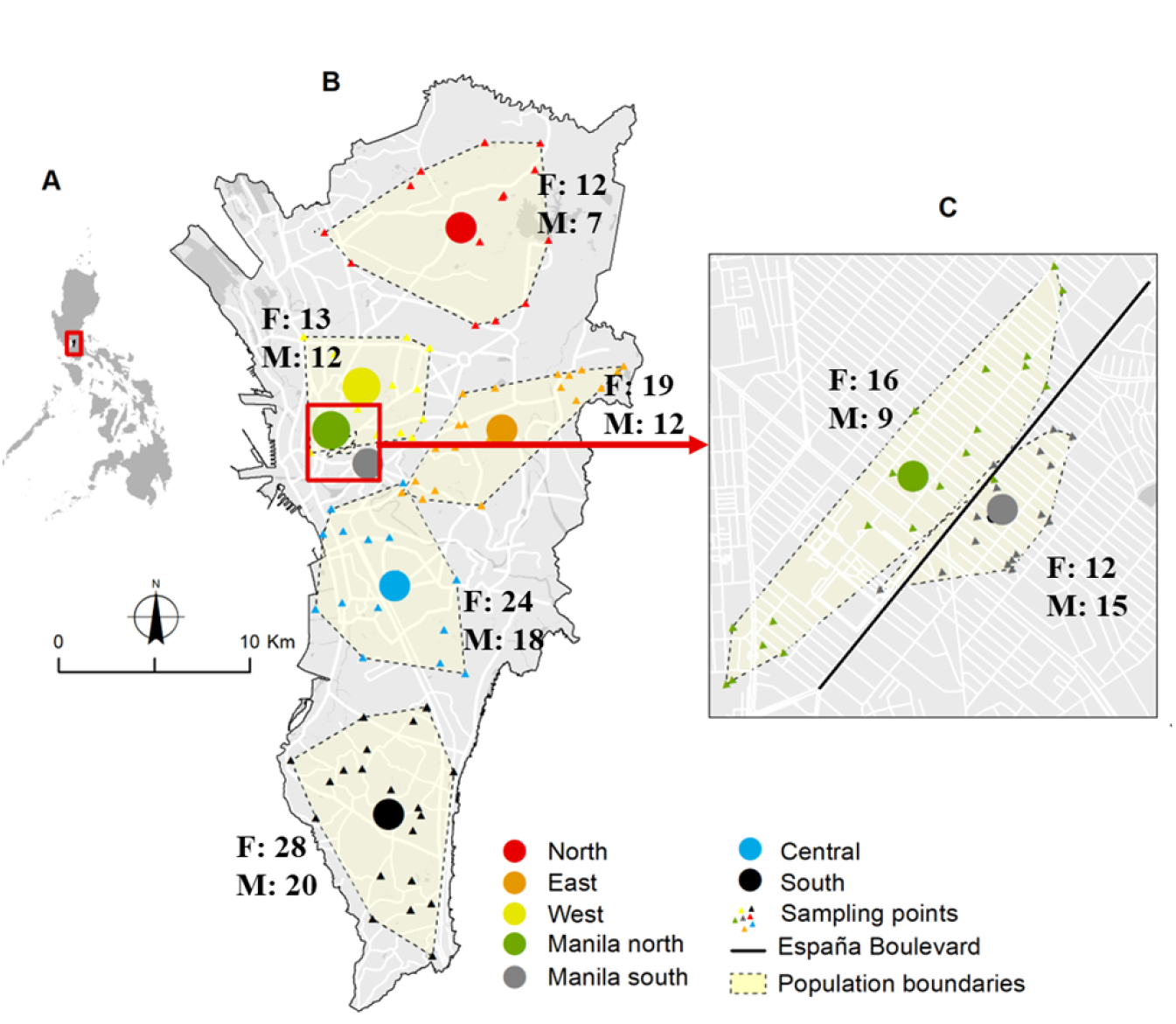
Geographic locations of *Ae. aegypti* collection sites in Manila City (C), Metropolitan Manila (B), Philippines (A). Circles indicate the geographical midpoints of *Ae. aegypti* populations per location; triangles indicate the households in the sampling locations. F and M indicate the total number of female and male individuals per population, respectively.

### DdRAD-Seq library preparation

Before ddRAD-Seq library preparation, the DNA concentration per individual was measured using a Quantus fluorometer (Promega, USA). We sequenced a pool of 7–28 individuals (S1Table) with equimolar DNA concentrations per individual (Pool-Seq) [31] in a population pool scheme.

To optimize library preparation, we compared seven restriction enzyme combinations (*DraI-NlaIII*, *MluCI-NlaIII* [19], *DraI-MluCI*, *SbfI HF-MspI* [20], *EcoRI-NlaIII* [19], *SbfI HF-HaeIII* [32], and *SspI-NlaIII*) to produce desired sequenceable DNA fragments of *c*.*a*. 100–500 bp, which following the addition of adapters and sequencing primers will result in an acceptable library size for sequencing (*c*.*a*. 200–700 bp). During optimization, we employed *in silico* and empirical digestion methods. Double digestion with *in silico* analysis allows the prediction of the number of sequenceable DNA fragments using restriction-site information from the enzymes and reference genome of *Ae. aegypti* [33] via the DDsilico program [19]. Empirical digestion analysis is an experimental method for observing DNA fragment distribution using the actual DNA of *Ae. aegypti* and restriction enzymes visualized with a High-Sensitivity DNA Assay 2100 Bioanalyzer (Agilent, USA). We selected *MluCI* and *NlaIII* (New England Biolabs, Beverly MA, USA) as the optimal combination because it generated the highest number of potential ddRAD loci using the *in silico* and empirical approaches (**S1 and S2 Figs.**). Rašić et al. [19] compared six different combinations of restriction enzymes, i.e., *SbfI-MseI*, *SphI-EcoRI*, *SphI-MluCI*, *SphI-MspI*, *NlaIII-EcoRI*, and *NlaIII-MluCI*, using *in silico* digestion alone for *Ae. aegypti*, finding that *MluCI* and *NlaIII* was the best combination. Because Rašić et al. [19] did not conduct an empirical digestion comparison, the validity of their finding complementarily supports the present study.

The DNA of each population pool was digested using the selected restriction enzymes (*MluCI* and *NlaIII*) for 3 h at 37°C. Subsequently, an enzyme inactivation step was performed by maintaining the DNA at 65°C for 20 min, after which a QiaQuick PCR Purification Kit (Qiagen, Hilden, Germany) was used to avoid further restriction enzyme activity in the samples. The digested DNA was then ligated to the modified Illumina P1 and P2 adapters [19]. Adapter ligation was performed using T4 Ligase buffer containing 0.5 µl of 4 nM/µl P1 Adapter, 0.5 µl of 6 nM/µl P2 Adapter, T4 DNA ligase, and H_2_O at 16°C for 16 h, after which the remaining ligase enzymes were inactivated at 65°C for 20 min. To increase the concentration of the adapter-ligated DNA (library), we amplified the library using PCR with a 10 µl reaction mixture containing 5 µl of Phusion High Fidelity MASTER Mix (New England Biolabs), 2 µl of P1 primer (AAT GAT ACG GCG ACC ACC GAG ATC TAC ACT CTT TCC CTA CAC GAC G), and 2 µl of P2 primer (CAA GCA GAA GAC GGC ATA CGA GAT CGT GAT GTG ACT GGA GTT CAG ACG TGT GC). The PCR conditions were as follows: 98°C for 30 s; 12 cycles of 98°C for 10 s, 60°C for 30 s, and 72°C for 90 s; and elongation at 72°C for 5 min. Seven PCR replicates were pooled and purified using a QiaQuick PCR Purification Kit to form the final library. Subsequently, the final library was checked for quality and quantity using the Bioanalyzer and a KAPA Quantification Kit (Roche, USA), respectively. The libraries were sequenced using a HiSeq X Ten Illumina sequencer (paired-end, 2 × 150 bp) at Beijing Genomics Institute, China.

### Data processing

The raw sequence data were assessed for sequence read quality using FASTQC v0.11.8 [34], and FASTQC information was used as a guide for trimming and filtering the raw data. Adapters and barcodes were removed using Trimmomatics 0.39 [35], and the trimmed and filtered data were mapped to the *Ae. aegypti* reference genome AaegL5.0 (www.vectorbase.org/organisms/aedes-aegypti/liverpool-lvp/AaegL5.0) using the *bwa mem* algorithm of the Burrows–Wheeler Aligner (BWA) [36]. The mapping results from BWA generated a mapping file in the Sequence Alignment Map (SAM) format. We filtered out ambiguously mapped reads with a minimum Phred quality score of <20. The SAM files were converted and sorted to Binary Alignment Map (BAM) files using SAMTOOLS 1.9 [37] to sort the sequences to the reference coordinates in a memory-efficient file form. All sorted BAM files of all populations were synchronized to the reference genome in the *mpileup* format using SAMTOOLS. The *mpileup* format file was converted to a *sync* file using java *mpileup2sync.jar* script on Popoolation2 [38]. SNPs were identified and allele frequencies estimated using *snp-frequency-diff.pl* script from Popoolation2 with a coverage range of 15–200 and minimum allele count of 4. Popoolation2 was used to perform SNP calls to reference genomes as reference call SNPs (rc SNPs) and SNPs observed between populations (pop SNPs), respectively. The rc SNPs were deleted, whereas the pop SNPs were retained for subsequent analysis.

### Detection of non-neutral SNP loci

We detected non-neutral SNP loci using three methods: empirical-, principal component analysis (PCA)-, and Bayesian-based methods. In the empirical-based method, we extracted non-neutral loci in the lower and upper 1% tails of a frequency distribution of pairwise *F*_ST_ values, as estimated via Popoolation2 using the *fst-sliding.pl* script with a coverage range of 15–200 and a minimum allele count of 4. SNPs detected in the lower tail were considered balancing selection candidates, whereas SNPs detected in the upper tail were considered divergent selection candidates [39]. The PCADAPT v.4.3.3 package in RStudio was employed to analyze Pool-Seq data [40]. In this analysis, the Benjamini–Hochberg procedure was used to decrease the false discovery rate during non-neutral loci detection [41]. The Bayesian-based method was employed using BayeScan 2.1 [42]. The input file used for BayeScan 2.1 was in the *bayenv* file format; therefore, we created a GenePop file from the *sync* file using the *subsample_sync2GenePop.pl* script in Popoolation2. The GenePop file was then edited and converted into the *bayenv* format using PGDSpider 2.1.1.5 [43]. BayeScan was run with 20 pilot runs, an additional burn in value of 50,000, and a thinning interval of 10. To further reduce false positive non-neutral loci detection, we defined non-neutral loci as those detected by at three methods. Subsequently, the detected non-neutral loci were removed from the complete dataset, and only the neutral SNP loci dataset was retained for population structure analysis. The non-neutral loci distribution was visualized across chromosomes using PhenoGram Plot (http://visualization.ritchielab.org/phenograms/plot) with some modification.

### Genetic diversity and population genetic structure

Genetic diversity per population was calculated using NPSTAT v.1.0 [44] by estimating the population mutation rate (Watterson’s ϴ) and nucleotide diversity (Tajima’s π) from a *pileup* file using a minimum Phred score of 20, coverage range of 15–200, and minor allele count of 4. Population genetic structure was analyzed using mean pairwise *F*_ST_ values across neutral SNP loci by generating an unweighted pair group method with arithmetic mean (UPGMA) dendrogram using Phylip-3.698 [45] and via nonmetric multidimensional scaling (NMDS) and permutational multivariate analysis of variance (PERMANOVA) analyses in the R package VEGAN v. 2.5-7 [46]. Mean pairwise *F*_ST_ values from neutral loci were also used to test isolation by distance for female and male populations by conducting the Mantel test via GenAlex 6.5 [47] and using a geographical distance matrix (km). Geographical distance was calculated based on the geographical midpoints of each population. The global *F*_ST_ values of female and male populations were calculated and then tested separately for neutral and non-neutral loci using the Wilcoxon rank-sum test in RStudio.

### Microsatellite analyses and comparing microsatellite markers with neutral SNP loci

To enable a comparison with the neutral SNP data obtained in the present study, we calculated the pairwise *F*_ST_ values among the 14 populations using previously obtained genotype data of 11 microsatellite markers at a regional scale [26] and a smaller spatial scale in Manila [27]. However, because different capillary electrophoresis instruments were used for fragment analysis in the previous studies [26,27], we separated the comparative analysis of 10 [26] and 4 [27] populations, respectively, to avoid bias due to instrumental differences. Microsatellite data were analyzed using ARLEQUIN v. 3.5 [48] to calculate pairwise *F*_ST_ values. The UPGMA dendrograms from the microsatellites and neutral SNPs of the 10 populations [26] were generated using Phylip-3.698 [45]. Additionally, a Mantel test was performed via GenAlex 6.5 [47] to test the correlation of pairwise *F*_ST_ values between microsatellite and neutral SNP markers in these 10 populations [26].

### Environmental association and gene annotation analyses of non-neutral SNPs

The non-neutral loci were used for environmental association analysis in which the environmental variables consisted of five climatic and eight landscape variables. We used the mean values of climatic variables (i.e., precipitation, air temperature, relative humidity, northward wind, and eastward wind) per population according to satellite-based remote sensing data obtained from the Google Earth Engine code editor platform [49] with an identical data duration and sampling collection time (**S1 Table**). Additional preprocessing to fill missing pixel data was performed using GRASS GIS software version 7.8.3 (GRASS Development Team, 2020). We used landscape data published by Francisco et al. [50], which included the percentage of the area of the following landscape categories in each village in 2014–2015 and 2017: water bodies, grassland, agricultural, open spaces, parks and recreation, residential areas, forests, and man-made buildings (education, health, and welfare; religious and cemetery; military; governmental institutions; industrial; commercial; transport; and informal settlements).

To assess the association between changes in allele frequencies in non-neutral loci and environmental conditions, we conducted distance-based redundancy analysis (db-RDA) using the *capscale* function and the variable selection algorithm via the *ordistep* function in the R package VEGAN v. 2.5-7 [46]. We used all climatic and landscape variables as explanatory variables and the pairwise genetic differences (*F*_ST_) matrix of each non-neutral locus as a response variable.

The non-neutral loci were annotated for candidate genes using BLASTX from the NCBI with its default parameters. The biological functions of the identified genes were investigated based on 259 categories of Gene Ontology annotations in the Universal Protein Knowledgebase (www.uniprot.org).

### Linkage disequilibrium of SNP loci

To assess linkage disequilibrium between SNP loci, we employed GENEPOP (genepop.curtin.edu.au) [51,52] with its default parameters, including a dememorization number of 1000, batch number of 100, and iterations per batch number of 1000. First, using the *subsample_sync2GenePop.pl* script in Popoolation2, we extracted the GenePop files of non-neutral SNP loci, separated by chromosome. Subsequently, the linkage disequilibrium between non-neutral SNP loci was tested for each chromosome. Focusing on clusters of non-neutral SNP loci in linkage disequilibrium with each other, we tested for linkage disequilibrium between the non-neutral SNP loci and the neutral SNP loci located between them.

## Results

### DdRAD-Seq and non-neutral SNP loci detection

In total, 377,047,648 raw reads with a length of 150 bp were generated via ddRAD-Seq, with 18,059,405 - 40,849,752 reads obtained per population. After trimming the adapters and filtering the low quality reads, 38,181,713 reads were removed and 338,865,935 reads were retained. In total, forward and reverse reads were covered 45.5 % of the entire reference genome (1,278,715,314 bp). We identified 65,473 SNP loci, with 1,880 and 2,401 SNP loci found exclusively among female and male populations, respectively.

The empirical-based method, PCADAPT, and BayeScan detected 655, 3,185, and 2,125 non-neutral SNP loci, respectively. Overall, 76 non-neutral SNP loci were detected by all three non-neutral detection methods (**S4 Fig.**). In subsequent analyses, the 76 non-neutral SNP loci detected by three non-neutral detection methods were classified as non-neutral SNP loci, whereas the other SNP loci were classified as neutral SNP loci. BLASTX searches found that 49 non-neutral SNP loci were located on or close to genes associated with enzymatic activity, nucleic acid-binding, and metabolism activity (**S4 Table**).

### Genetic diversity and population genetic structure

Watterson’s ϴ and Tajima’s π of neutral and non-neutral loci indicated low genetic diversity throughout all populations (Watterson’s ϴ = 0.011-0.012; Tajima’s π = 0.010-0.011). Mean pairwise *F*_ST_ values among the populations were 0.022 – 0.069 (overall mean = 0.038) for neutral loci and 0.042-0.23 (overall mean = 0.120) for non-neutral loci. Mean global *F*_ST_ values were compared across neutral SNP loci using a Wilcoxon rank-sum test; significant differences were found between female and male populations (*p* < 0.05), but no significant differences were found across non-neutral loci (*p* > 0.05). PERMANOVA of the neutral loci indicated that no significant difference existed between female and male populations, and NMDS plots revealed an unclear separation between the female and male populations based on neutral loci (**Fig. 2**). Male populations exhibited a more diverse structure than that of female populations, with the North Manila and South Manila populations being genetically isolated from the other populations.

**Fig 2.**
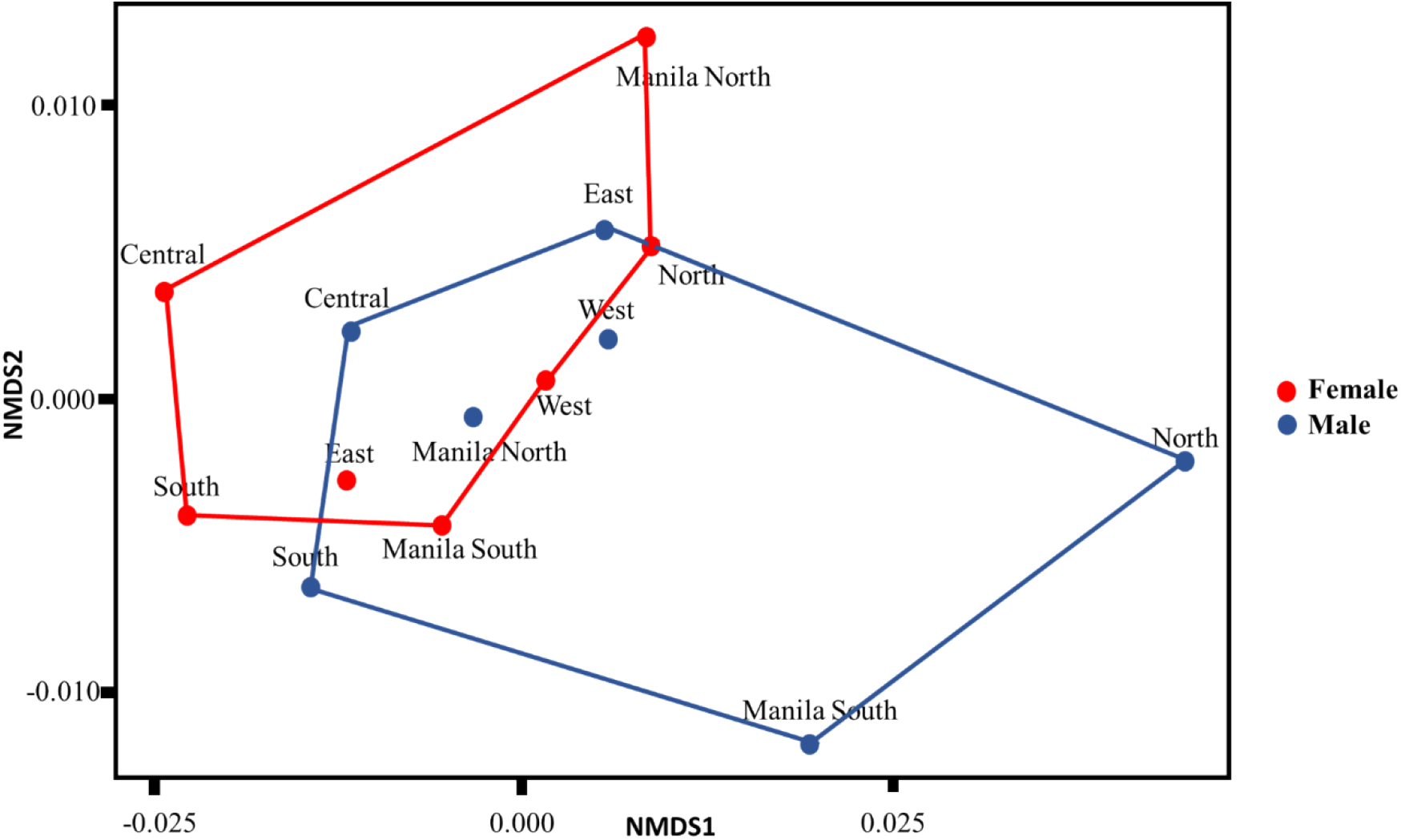
Population genetic structure of male (blue) and female (red) *Ae. aegypti* populations according to nonmetric multidimensional scaling (NMDS) based on neutral loci. PERMANOVA analysis revealed no significant divergence between the female and male populations (R^2^ = 0.09141, *p* = 0.2366).

### Comparison of neutral SNP and microsatellite markers

In the 10 studied populations in Metropolitan Manila, the pairwise *F*_ST_ values generated from neutral SNP loci (0.022 - 0.069; mean = 0.039) were higher than those generated from microsatellite markers (0–0.043; mean = 0.015). According to the Mantel test, the pairwise *F*_ST_ values obtained from neutral SNP loci and microsatellite markers were significantly correlated (p < 0.05; Fig. 3) with a regression equation of y = 0.3327x + 0.0336.

**Fig 3.**
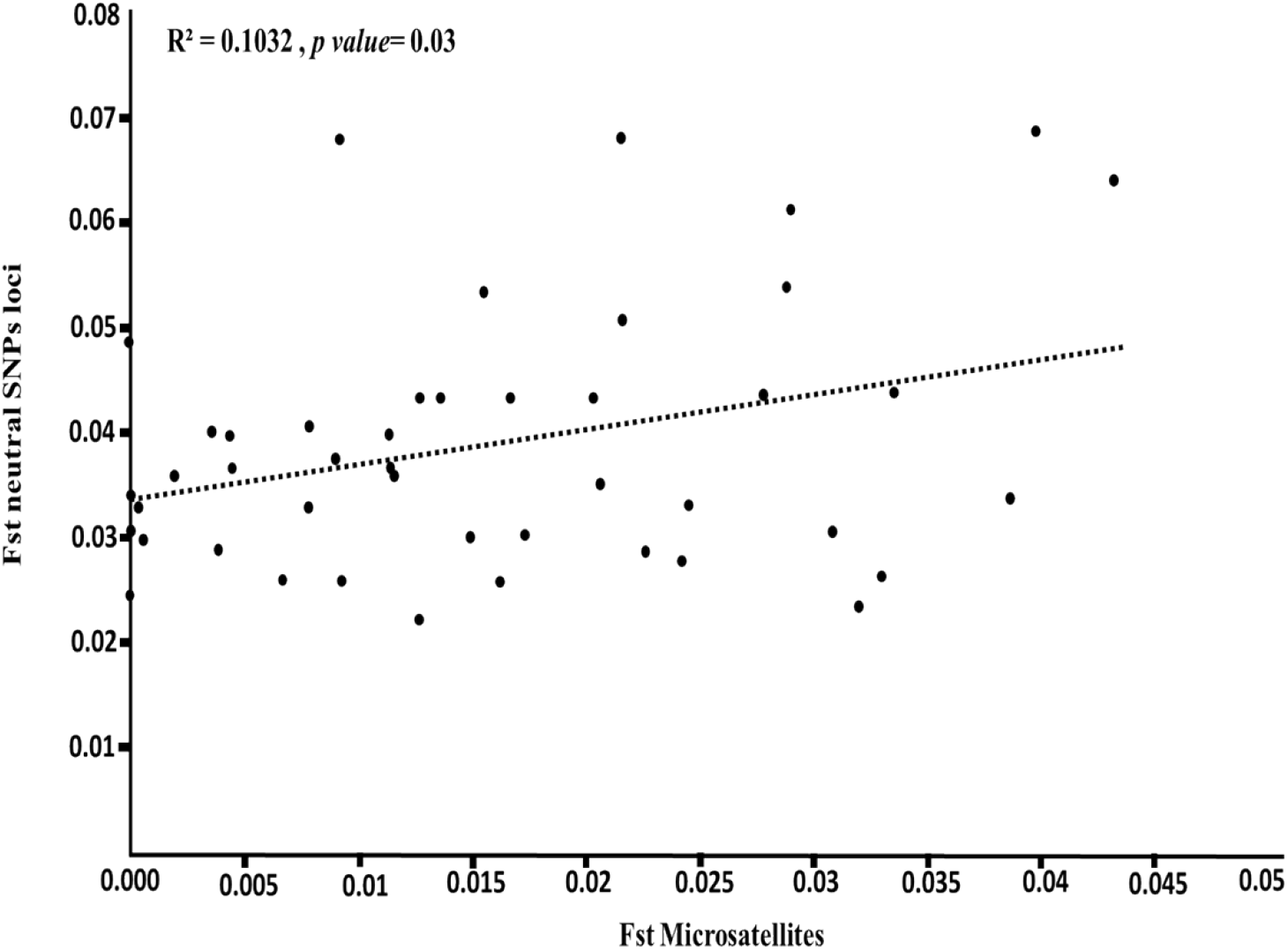
Regression of pairwise genetic differentiation (*F*_ST_) of neutral SNP and microsatellite markers obtained using data from Metropolitan Manila.

Figure 5 compares the dendrograms of the 10 *Ae. aegypti* populations in Metropolitan Manila constructed using the pairwise *F*_ST_ values of microsatellite and neutral SNP markers. The dendrogram generated from microsatellite markers had shallow branch lengths (0.010) from the terminals to the common ancestor. In contrast, in the dendrogram generated from neutral SNP loci, the overall branch length from the terminal to the common ancestor was longer (0.030), indicating the unique genetic structure of individual populations. Additionally, based on the dendrogram of microsatellites markers (Fig. 4b), eight populations (Female North with Male North, Female West with Male West, Female East with Male East, and Female South with Male Central) were not separated in the dendrogram. Besides that, by utilizing neutral SNPs loci, almost all pairs of populations were separated from each other. At a smaller scale, i.e., within Manila, the *F*_ST_ values of the neutral SNP loci of Female North–South Manila and Male North–South Manila were slightly higher (0.037 and 0.040, respectively) than those of microsatellite markers (0.037 and 0.024, respectively).

**Fig 4.**
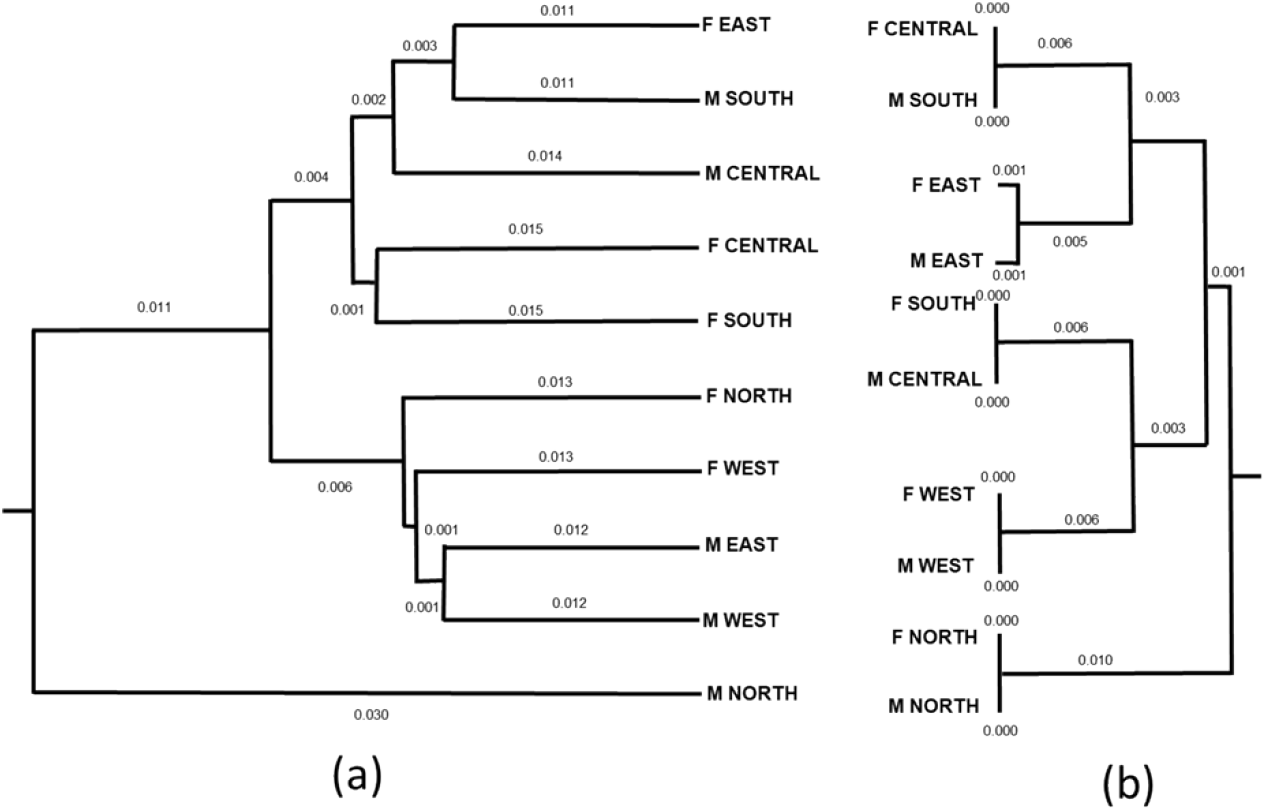
“Unweighted pair group method with arithmetic mean” dendrograms based on mean pairwise genetic differences (*F*_ST_) among 10 regional-scale populations in Metropolitan Manila according to (a) neutral SNP loci and (b) microsatellite markers.

**Fig 5.**
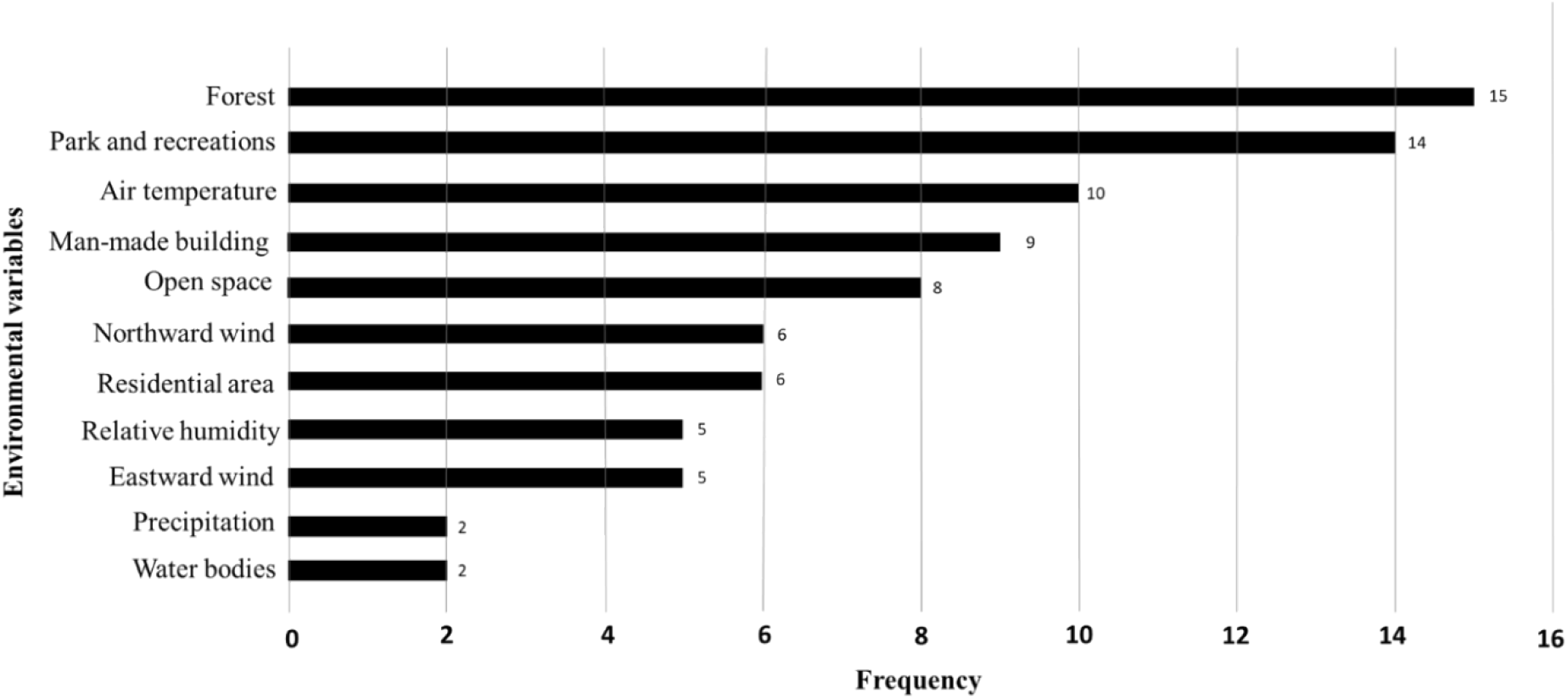
Frequency of 57 non-neutral loci associated with environmental variables selected in distance-based redundancy analysis models.

### Association of non-neutral SNP loci and environmental variables

Db-RDA analysis revealed that 57 of 76 non-neutral SNP loci were significantly associated with 12 environmental variables (Fig. 5). Of these non-neutral SNP loci, 36, 12, and 9 were associated with landscape variables, climatic variables, and both landscape and climatic variables, respectively. Forest was associated with the highest number of non-neutral SNP loci (15), followed by park and recreation (14), air temperature (10), man-made building (9), and open space (8). The other 6 environmental variables were associated with 2–6 non-neutral SNP loci.

Linkage disequilibrium tests revealed that all non-neutral SNP loci were in linkage disequilibrium (*p* < 0.05) within each of the four clusters of physically close (i.e., 2–41 bases) non-neutral SNP loci on a chromosome (Fig. 6). Additionally, db-RDA analysis revealed that non-neutral SNP loci within the same cluster were associated with the same environmental variables. Neutral SNP loci located in clusters of non-neutral SNP loci did not show linkage disequilibrium with each other (*p* > 0.05), whereas non-neutral SNP loci always exhibited linkage disequilibrium (*p* < 0.05) with another non-neutral SNP loci in one cluster (**S3 Table**).

**Fig. 6.**
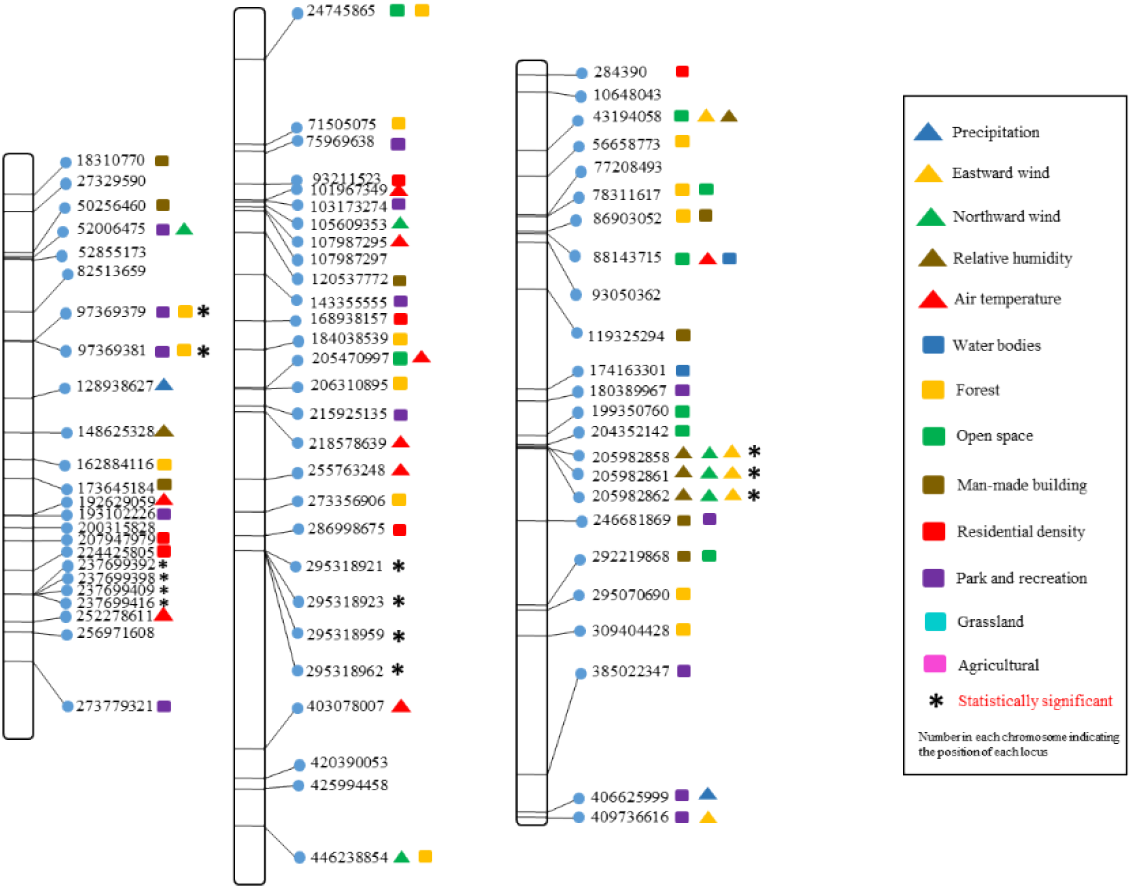
Map of 76 non-neutral SNP loci across three chromosomes of *Ae. aegypti*. Colored symbols indicate the environmental variables to which the non-neutral SNP loci are associated. The (*) indicate statistically significant (*p-value* < 0.05) with Fisher’s method in GENEPOP.

## Discussion

### Adaptive divergence of *Ae. aegypti* populations

In this study, 76 non-neutral SNP loci were detected in *Ae. aegypti* populations at a fine scale, i.e., in Metropolitan Manila, and more of these non-neutral SNP loci (n = 36) were associated with landscape variables than climatic variables (n = 12) (Fig. 5), which may have been due to the spatially homogeneous climatic conditions across Metropolitan Manila, despite the high genetic divergence among the 14 studied populations. It is usually difficult to measure the spatial heterogeneity of microclimates at an intraurban scale, e.g., in residential landscapes, in which microclimates are homogenous [53,54].

Of the 57 non-neutral SNP loci associated with landscape and climatic variables, 39 were associated with forest, parks and recreation, air temperature, man-made building and open space (Fig. 5) which are consistent with the result of [13] showing the genetic variation of SNPs was correlated with vegetation and temperature. Such areas are known to be associated with reproduction- and survival-related fitness in *Ae. aegypti*, which is abundant in urban green spaces [55,56]. Mosquito larvae live in water habitats, including man-made containers, drains, canals, and ponds [57,58]. In addition, both male and female mosquitoes require vegetation as a sugar source and resting place to aid blood ingestion [59]. Vegetation also supplies local moisture, supporting the activity and survival rate of mosquitoes [60,61]. In urban areas, medically important mosquitoes, such as *Ae. aegypti* and *Ae. albopictus*, are found in parks and green spaces [56,62,63]. According to evolutionary theory, fitness and adaptive divergence are related [64,65], i.e., high fitness or suitability to a specific environmental condition may increase the local density of a species, leading to high intraspecific competition for space or resources. Consequently, some populations adaptively evolve to alternative environmental conditions to avoid competition, leading to adaptive divergence along environmental gradients [66]. Although we did not measure mosquito abundance or the level of intraspecific competitive pressure in the present study, we found that environmental factors potentially cause adaptive divergence at many non-neutral SNP loci, which is associated with mosquito fitness and supports the aforementioned theory.

The detected non-neutral SNP loci were physically clustered in several positions across three *Ae. aegypti* chromosomes and exhibited linkage disequilibrium (Fig. 6). Loci in physical proximity are more likely to be in linkage disequilibrium than loci located at distant positions on the same or other chromosomes because they are less likely to cross over during recombination [67]. We also found that non-neutral and neutral SNP loci within physically close clusters exhibited and did not exhibit linkage disequilibrium, respectively. A study on Japanese cedar, *Cryptomeria japonica* [68], found that the frequency of linkage disequilibrium among non-neutral loci was relatively high because these loci tended to be closely associated. Interestingly, we found that non-neutral SNP loci in linkage disequilibrium within the same cluster were associated with a common environmental variable (**S3 Table**). Non-neutral SNP loci are presumed to be affected by directional or balancing selection. If one physically close and linked non-neutral SNP locus undergoes natural selection due to an environmental variable, the other non-neutral SNP locus may exhibit allele frequency changes similar to those of the naturally selected locus, even if it has no genetic function or its genetic function is unrelated to the natural selection. Therefore, our results do not necessarily indicate that groups of non-neutral loci with similar functions are located in physical proximity on chromosomes.

### Contrasting spatial genetic structures of male and female *Ae. aegypti* populations

The spatial population genetic structures of female and male *Ae. aegypti* exhibited contrasting patterns according to neutral SNP loci, i.e., male populations exhibited greater spatial divergence than female populations (Fig. 2). Medeiros et al. [69] previously found that the dispersal ability of male and female mosquitoes was different. This pattern was also consistent with another study in Metropolitan Manila [29], in which wing geometry and microsatellite markers were employed. The different dispersal abilities of male and female mosquitoes might be due to differences in their behavior, such as sex-specific feeding preference and host-seeking behavior [29,63]. In addition, higher genetic variation in male populations suggests gene flow may be low between male populations, reflecting their low dispersal ability [70]. Maciel-De-Freitas et al. [71] investigated the dispersal and survival rates of *Ae. aegypti* in Rio de Janeiro, finding that female mosquitoes move farther than males (mean flight distance: females = 40.94–78.81 m; males = 32.02–42.26 m) and tend to live longer than males. Female *Ae. Aegypti* mosquitoes can fly long distances in search of blood meals or oviposition sites when these resources are not available within close range. Indeed, gravid female *Ae. aegypti* can fly up to several kilometers to find a suitable egg laying site [70]. Furthermore, female *Ae. aegypti* undertake more interactions with humans than males, giving them more opportunity to experience long-distance passive dispersion via human transportation networks [29], which would contribute to increasing the flight distance of female mosquitoes during host-seeking behavior. However, in this explanation of the different dispersal abilities of male and female mosquitoes, the different number of SNPs found in each sex is not considered. Indeed, both sex differences in dispersal ability and the higher number of polymorphic loci in the male genome may have contributed to the high spatial genetic divergence detected in males.

### Advantages and limitations of the pooled ddRAD-Seq approach: comparison with microsatellite markers

We found a significant positive correlation between the pairwise *F*_ST_ values of neutral SNP loci and microsatellite markers (Fig. 3), which is consistent with previous studies on Atlantic salmon, *Arabidopsis halleri*, Gunnison sage-grouse, corn rootworm, and alpine-endemic birds [72–79]. In addition, we found that the pairwise *F*_ST_ values of neutral SNP markers were higher than those of microsatellite markers, indicating higher resolution in terms of measuring population differentiation. In Fig. 3, the intercept of the regression line (0.0336) indicates that neutral SNP markers can detect a certain amount of genetic differentiation (i.e., pairwise *F*_ST_ = 0.0336), even among populations in which microsatellite markers do not detect any genetic differentiation.

Genetic variation determined at a small spatial scale, such as in Metropolitan Manila, tends to be low (e.g., isolation by distance [80]). However, ddRAD-Seq can detect a large number of SNP loci, increasing sensitivity for the detection of low genetic variation that cannot be detected with microsatellite markers. Previous studies [81,92] revealed that, at a fine-spatial scale, microsatellite markers perform better than a small number of SNP markers for determining population differences because of their high mutation rate. Although microsatellite markers are highly polymorphic, only a small number are usually used owing to logistical constraints [15]. In contrast, Ryynänen et al. [72] found that a small number of SNP markers and microsatellite markers provided comparable results. At a fine-spatial scale, the genetic differences among individuals in different populations are difficult to distinguish because they can share high kinship owing to their adjacent habitat. To determine genetic variation between populations on a small spatial scale, a high number of SNP loci should be used to detect population structure [72,74]. A few thousand SNPs identified via NGS, compared with several microsatellite markers, are sufficient to estimate the genetic diversity and divergence in natural populations [74,75].

We applied the Pool-Seq approach with ddRAD-Seq. In recent years, this approach has increasingly be used for population genomic studies [74,83–90]. When taking an individual-based approach, only a limited number of individuals are selected for analysis from a population owing to cost constraints, and sampling errors that occur when selecting these individuals can lead to erroneous allele frequency estimates in the population. In contrast, in Pool-Seq, libraries are constructed per population rather than per individual, which markedly reduces the resources (e.g., reagents) and time required to complete the analysis. Thus, many more individuals per population can be analyzed, reducing the sampling error and increasing the accuracy of allele frequency estimation [31]. Notably, the Pool-Seq approach has several limitations. First, it cannot recognize the haplotypes of each individual; thus, some analyses, such as STRUCTURE [91] (a widely used individual-based population genetic analysis method), cannot be performed with Pool-Seq. Second, if the amount of template DNA for each individual is unequal when pooled samples are prepared, heterogeneity increases substantially during PCR amplification and may reduce the accuracy of allele frequency estimation [31]. Therefore, in a population pool, we mixed the same amount of DNA per individual in a population.

In conclusion, we successfully used a pooled ddRAD-Seq approach to detect adaptive divergence among *Ae. aegypti* populations along environmental gradients occurring at a relatively fine-spatial scale in Metropolitan Manila. Additionally, we detected dispersal patterns among local populations and their sex differences using this approach. Non-neutral loci analysis revealed that spatial heterogeneity in landscape factors related to mosquito fitness may cause adaptive divergence in the *Ae. aegypti* populations of Metropolitan Manila. Furthermore, data on vector mosquito dispersal at a fine-spatial scale could be used to design and impellent vector control programs, such as *Wolbachia*-infected mosquito mass-release programs, which will benefit from information on mosquito population dispersal patterns and the potential barriers to mosquito movement in and around the release area.

## Acknowledgments

The authors would like to thank Dr. Belinda Kahnt (Martin-Luther-Universität Halle-Wittenberg, Institute of Biology, General Zoology) for assisting with the bioinformatics work, Micanaldo Ernesto Francisco (Ehime University) for helping make the map of Metropolitan Manila and providing the environmental data of Metropolitan Manila, and Dr. Maribet Gamboa (Universidad Catolica de la Santisima Concepcion) for improving our understanding of ddRAD-Seq.

